# *Listi*Wiki: a database for the foodborne pathogen *Listeria monocytogenes*

**DOI:** 10.1101/2023.10.16.562455

**Authors:** Christoph Elfmann, Bingyao Zhu, Jörg Stülke, Sven Halbedel

## Abstract

*Listeria monocytogenes* is a Gram positive foodborne pathogen that regularly causes outbreaks of systemic infectious diseases. The bacterium maintains a facultative intracellular lifestyle; it thrives under a variety of environmental conditions and is able to infect human host cells. *L. monocytogenes* is genetically tractable and therefore has become an attractive model system to study the mechanisms employed by facultative intracellular bacteria to invade eukaryotic host cells and to replicate in their cytoplasm. Besides its importance for basic research, *L. monocytogenes* also serves as a paradigmatic pathogen in genomic epidemiology, where the relative stability of its genome facilitates successful outbreak detection and elucidation of transmission chains in genomic pathogen surveillance systems. In both terms, it is necessary to keep the annotation of the *L. monocytogenes* genome up to date. Therefore, we have created the database *Listi*Wiki (http://listiwiki.uni-goettingen.de/) which stores comprehensive information on the widely used *L. monocytogenes* reference strain EDG-e. *Listi*Wiki is designed to collect information on genes, proteins and RNAs and their relevant functional characteristics, but also further information such as mutant phenotypes, available biological material, and publications. In its present form, *Listi*Wiki combines the most recent annotation of the EDG-e genome with published data on gene essentiality, gene expression and subcellular protein localization. *Listi*Wiki also predicts protein-protein interactions networks based on protein homology to *Bacillus subtilis* proteins, for which detailed interaction maps have been compiled in the sibling database *Subti*Wiki. Furthermore, crystallographic information of proteins is made accessible through integration of Protein Structure Database codes and AlphaFold structure predictions. *Listi*Wiki is an easy-to-use web interface that has been developed with a focus on an intuitive access to all information. Use of *Listi*Wiki is free of charge and its content can be edited by all members of the scientific community after registration. In our labs, *Listi*Wiki has already become an important and easy to use tool to quickly access genome annotation details that we can keep updated with advancing knowledge. It also might be useful to promote the comprehensive understanding of the physiology and virulence of an important human pathogen.

## INTRODUCTION

*Listeria monocytogenes* is a dreaded human pathogen causing foodborne infections that may develop into severe systemic disease in susceptible risk groups. Case fatality of such invasive infections approaches 30-40% (Charlier et al., 2017), which is exceptionally high compared to other gastrointestinal bacterial pathogens (Werber et al., 2013). The natural reservoir of the bacterium is the environment, where it lives in the soil and is found associated with plant surfaces, but also in fresh and marine water samples, from where it is carried over into food production (Quereda et al., 2021; Vivant et al., 2013). For systemic infection, the bacterium breaches anatomical barriers such as the gut epithelium or the blood brain barrier by repeated cycles of host cell invasion, escape from phagosomes, intracellular growth in the cytoplasm of infected host cells and spread to uninfected neighbor cells (Dussurget et al., 2004; Vazquez-Boland et al., 2001).

*L. monocytogenes* belongs to the *Bacillota* phylum (formerly known as firmicutes) and is the best-established model organism to study the mechanisms of bacterial growth inside mammalian host cells within this phylogenetic group (Hamon et al., 2006). The bacterium can switch between the contrasting lifestyles of a peaceful environmental soil dweller and a harmful intracellular pathogen. This ability has made *L. monocytogenes* a popular model system to study adaptation to stresses experienced in the environment, in the gastrointestinal tract or during confrontation with the immune system (Freitag et al., 2009; Leseigneur et al., 2020).

The reconstruction of the genome sequence of the widely used reference strain EGD-e in 2001 provided first insights into the genetic repertoire of the pathogen (Glaser et al., 2001). Global analyses using omics technologies further improved functional annotation of the *L. monocytogenes* genome. These analyses include the definition of compartment specific proteomes (Desvaux et al., 2010; Dumas et al., 2009; Ramnath et al., 2003), genome wide *in silico* prediction of subcellular protein localization (Renier et al., 2012) and the definition of transcriptomes under various conditions and in different strains (Behrens et al., 2014; Chatterjee et al., 2006; Hain et al., 2012). In 2014, few single nucleotide polymorphisms in the EGD-e genome were found in a genome resequencing project (Becavin et al., 2014), possibly reflecting ongoing microevolution in the lab. However, genomic analysis of *L. monocytogenes* isolates collected from patients, environmental and food sources in pathogen surveillance programs revealed an even higher diversity within the species that separates into at least 4 phylogenetic lineages, which even differ in their virulence gene content (Moura et al., 2016; Yin et al., 2019).

Beyond the use of genome-wide approaches, the annotation of the *L. monocytogenes* genome has been continuously improved by the identification of the function of previously uncharacterized genes. While the function of about 1,000 genes remained unclear in the original annotation (Glaser et al., 2001), only about 350 genes are currently without any further functional annotation and this number will further decrease.

To combine our knowledge of the genetic elements present in the *L. monocytogenes* EGD-e genome in a single database and to keep this annotation updated on a regular basis, we have designed *Listi*Wiki, an user-editable database that integrates the available information on all protein and RNA encoding genes, their transcriptional organization, functional assignments and the mutual interactions of their gene products based on the published body of knowledge. The *Listi*Wiki dataset should not only provide a comprehensive collection of genomic annotation details of the EGD-e reference strain, it also may offer an up-to-date reference for automated genome annotations of new *L. monocytogenes* isolates.

## DESCRIPTION OF THE DATABASE

*Listi*Wiki (listiwiki.uni-goettingen.de) uses the same framework as a series of other model organism databases developed by our group (*Subti*Wiki, *Syn*Wiki, and *Myco*Wiki) (Elfmann et al., 2022; Pedreira et al., 2022a; Pedreira et al., 2022b). Following the same general layout of the websites and the organization of data, the core of *Listi*Wiki’s structure is made up by the *Gene pages* for each individual gene. These pages are dedicated to the description of genes and the corresponding proteins. While most of the information relating to the gene in question can be found directly on the page, some aspects of the gene and the encoded product can be explored further using several interactive *browsers*, such as the *Genome, Interaction* and *Expression browsers*. Additional pages provide information about groups of genes (such as the *Categories page*) or offer more functionality to the user (such as the *Export page*).

### Gene identifiers

On *Listi*Wiki, different identifiers are used to designate genes. The *gene name* is usually a three-to-four letter mnemonic, using the standard nomenclature for bacterial genes (such as *gpsB*). The gene designations are derived from the most recent annotation for *L. monocytogenes* strain EGD-E (RefSeq accession: NC_003210.1) (Toledo-Arana et al., 2009). Moreover, we assigned 328 gene names based on similarity to *B. subtilis* genes. As the names should be ideally unique, manual curation was necessary in some cases to avoid duplicates. For example, the gene designation *prfA* is used twice in the RefSeq entry, once for the major transcriptional regulator of virulence genes in *L. monocytogenes*, and once for the conserved peptide chain release factor 1, which also exists in *L. monocytogenes*. Thus, to avoid confusion, we renamed the latter to *prf1* in agreement with the UniProt database. Another identifier is the *locus tag*, which is a unique label assigned to each genomic feature during genome annotation (Glaser et al., 2001; Toledo-Arana et al., 2009). For example, *lmo1888* is the locus tag of the *gpsB* gene. While the names of genes can change (*e*.*g*., when a new name is assigned to a previously uncharacterized gene), the locus tag will remain constant. For some genes, only the locus tag is provided, which is then used as the primary identifier of the gene and is displayed as the title of the corresponding Gene page.

### The Front page

The Front page gives access to many of *Listi*Wiki’s features (see Fig. 1). Its most important feature is the search bar, which allows to look up genes by different keywords and identifiers, but also performs full-text searches. In addition, the Front page displays various navigation elements. With the top bar, users can access the Categories page, which presents a tree-like structure of functional categories to which the genes and corresponding proteins of *L. monocytogenes* are assigned. Furthermore, the top bar gives access to a list of essential genes and contains buttons to access a random Gene page. Finally, it allows to log in or register for *Listi*Wiki.

**Figure 1.**
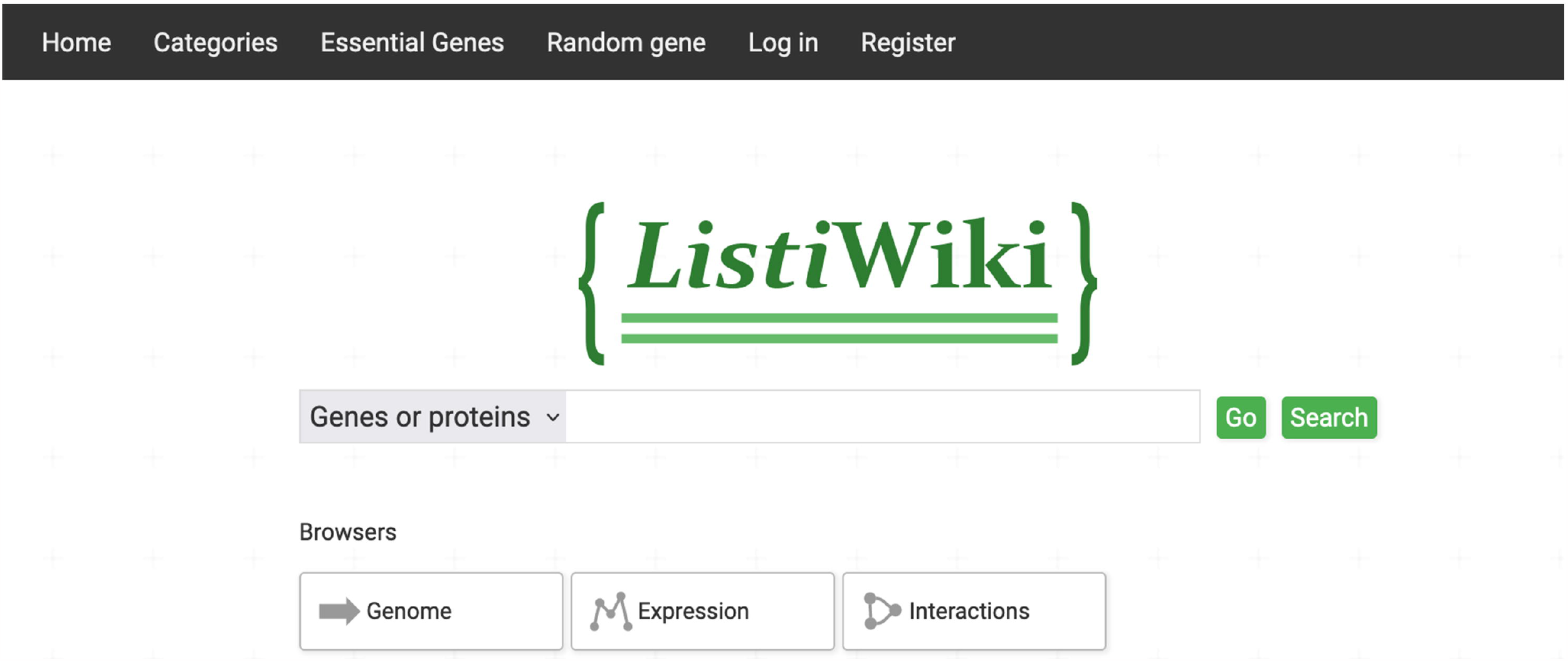
The Front page of *Listi*Wiki. The search bar in the center of the pages allows to search for genes by identifiers and keywords. The top bar gives access to an overview of categories, a list of essential genes, and a random gene page. It also provides functionality for login and registration. Below the search bar, the interactive browsers can be directly accessed.

### The Gene pages

The *Gene pages* represent the core of *Listi*Wiki and its underlying data structure. As mentioned above, most of the information pertaining to a specific gene and the encoded protein can be found on the corresponding Gene page. In addition, the user can explore the role of the gene in more detail by using several browsers, which can be accessed via the top navigation bar. The latter also features a search bar for convenience, as well as log-in functionality. Figure 2 displays the Gene page for *gpsB*.

**Figure 2.**
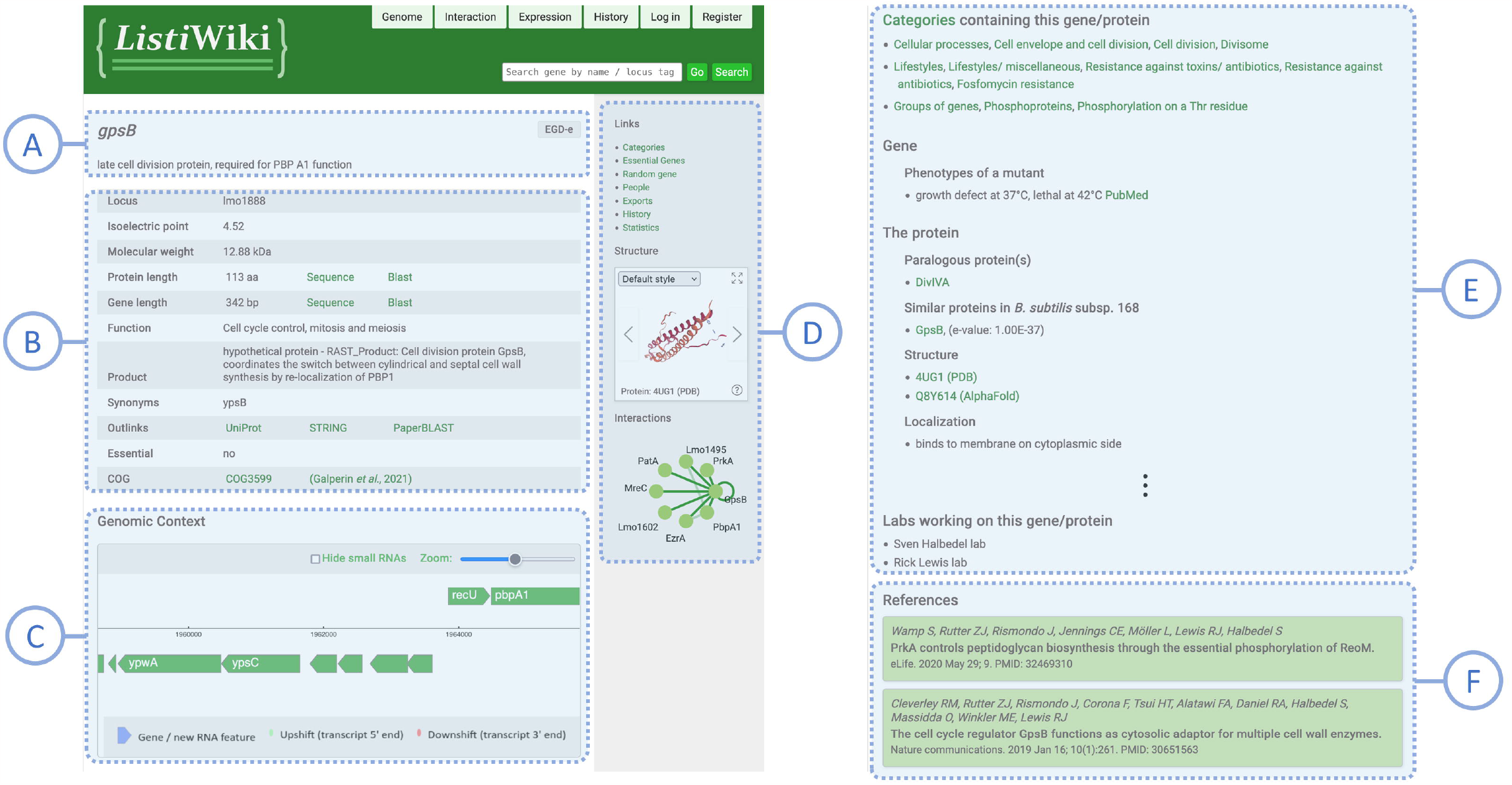
The Gene page for *gpsB*. Each Gene page has the same general structure, but depending on the available data, individual elements might vary. **(A)**, page header with gene name, strain and description; **(B)**, tabular overview over basic properties of the gene and its protein; **(C)**, Genomic Context Viewer, an interactive display showing the genomic neighborhood of the gene; **(D)**, side bar with navigation links, the Structure Viewer and the Interaction Graph; **(E)**, various sections on different aspects of the gene, such as assigned functional categories or the protein; **(F)**, list of relevant publications. Some sections were omitted in the figure as indicated by the vertical ellipsis.

While the data available for each gene may vary, the general structure of each Gene page is the same. The main body is introduced with the gene name as the page title, followed by a short general description. Next, a table summarizes basic properties of the gene, such as the locus tag, isoelectric point and molecular weight of the protein, details about the associated sequences and additional information about function and product. Utility links allow to view the associated sequences in the Genome browser, and to directly perform a BLAST search (Sayers et al., 2022) with them. The table also contains outlinks to the protein database UniProt (UniProt, 2023) the interaction database STRING (Szklarczyk et al., 2023) and PaperBLAST (Price and Arkin, 2017), a tool to find related publications based on a protein sequence. The last row displays the Cluster of Orthologous Genes (COG) (Galperin et al., 2015) the gene belongs to.

Below this tabular overview, the Genomic Context Viewer is embedded, an interactive display for the genomic neighborhood of the gene. The viewer is a simplified version of the interactive *Genome Browser*, which will be discussed below. Next, different sections of text describe the gene/protein in further detail. For example, most genes are assigned to several functional categories, which have been specifically curated for *Listi*Wiki. A list of these assigned categories can be found as one of the Gene page sections. Additional sections describe properties of the encoded protein or the regulation of the gene or operon among others, in more detail. Moreover, the Gene pages list biological materials and relevant publications.

In the side bar, several navigation links give access to other pages, such as the Categories page, or the Export page. The latter allows users to download tabular data on all entries of a certain entity, such as genes, categories or operons. The side bar also features more information on the gene or protein in the form of graphical displays. One such display is the interactive *Structure Viewer*, which displays three-dimensional protein structures (see Fig. 3A). The user can cycle through the different structures available for the protein in question, such as experimentally derived structures from PDB (Berman et al., 2000). The viewer also displays structure predictions from the AlphaFold Protein Structure Database (Varadi et al., 2022), which we added for all proteins in the *Listi*Wiki database. The Structure Viewer features a full-screen mode and different style options, for example coloring the protein surface based on hydrophobicity. The rendering of the molecules is performed with NGL Viewer (Rose and Hildebrand, 2015). Another feature of the sidebar is the Interaction Graph, an interactive display presenting protein-protein interactions (see Figure 3B). The user can click on the protein nodes to navigate to the corresponding Gene page or select edges to show further information on the corresponding interactions. A basic version of this graph was already implemented in *Subti*Wiki (Pedreira et al., 2022b), but we recently expanded its capabilities. Before, only the interactions between the protein in question and its direct interaction partners where shown, but now interactions *between* these partners are included, as well. The latter are displayed as additional transparent edges between the nodes.

**Figure 3.**
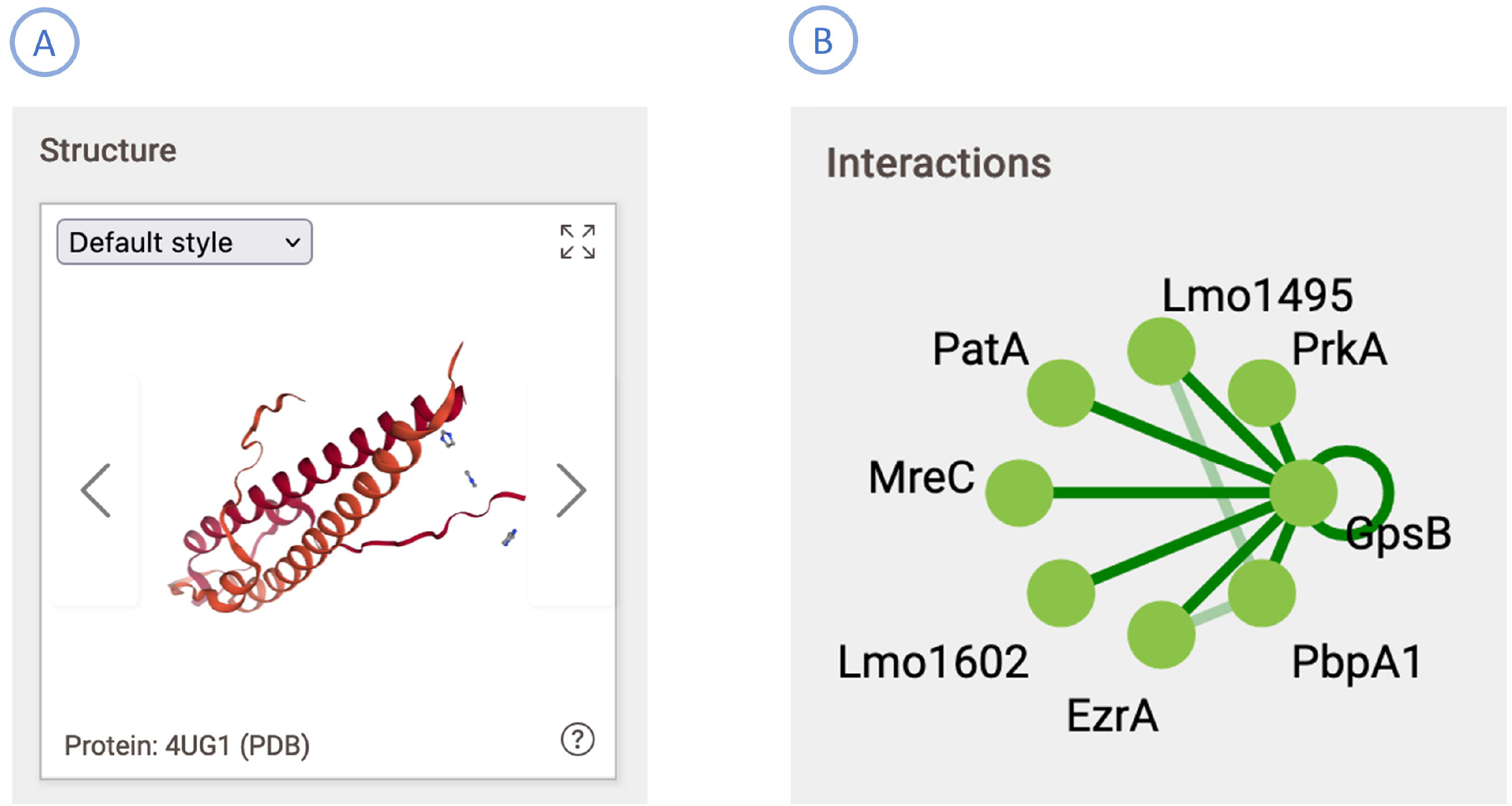
The interactive displays available in the side bar of the Gene page. **(A)** The Structure Viewer displays the structures available for a protein (here: GpsB). In addition to experimentally determined structures from PDB, structure predictions from the AlphaFold Protein Structure Database are presented, as well. Different rendering options and a full-screen mode are available. **(B)** Interaction Graph for GpsB showing its protein-protein interactions. Interactions of GpsB are displayed as opaque edges, while interactions between the other interaction partners are transparent.

### The Browsers

Using the top bar of any Gene page, the user can access several browsers, which are pages dedicated to specialized interactive tools. They allow users to explore certain aspects of the organism’s biology in more detail. When accessed from the top bar of a Gene page, the data corresponding to that gene is highlighted in the selected browser. But each browser also features its own search bar, which makes it easier to access and compare information on different genes or proteins. Currently, three browsers are featured on *Listi*Wiki.

The Genome Browser (Fig. 4) allows the user to view the genomic properties of the gene in question, such as its coordinates and orientation, as well as its nucleotide and amino acid sequence. The featured interactive graphical display also shows the neighborhood of the gene, which the user can scroll through. By clicking on any gene marker in the display, the associated sequences are loaded. The user can then search the sequences for substrings and toggle the reverse complement. The tabs at the top of the page can be used to perform advanced searches, such as querying arbitrary regions of the genome or genes including their flanking regions.

**Figure 4.**
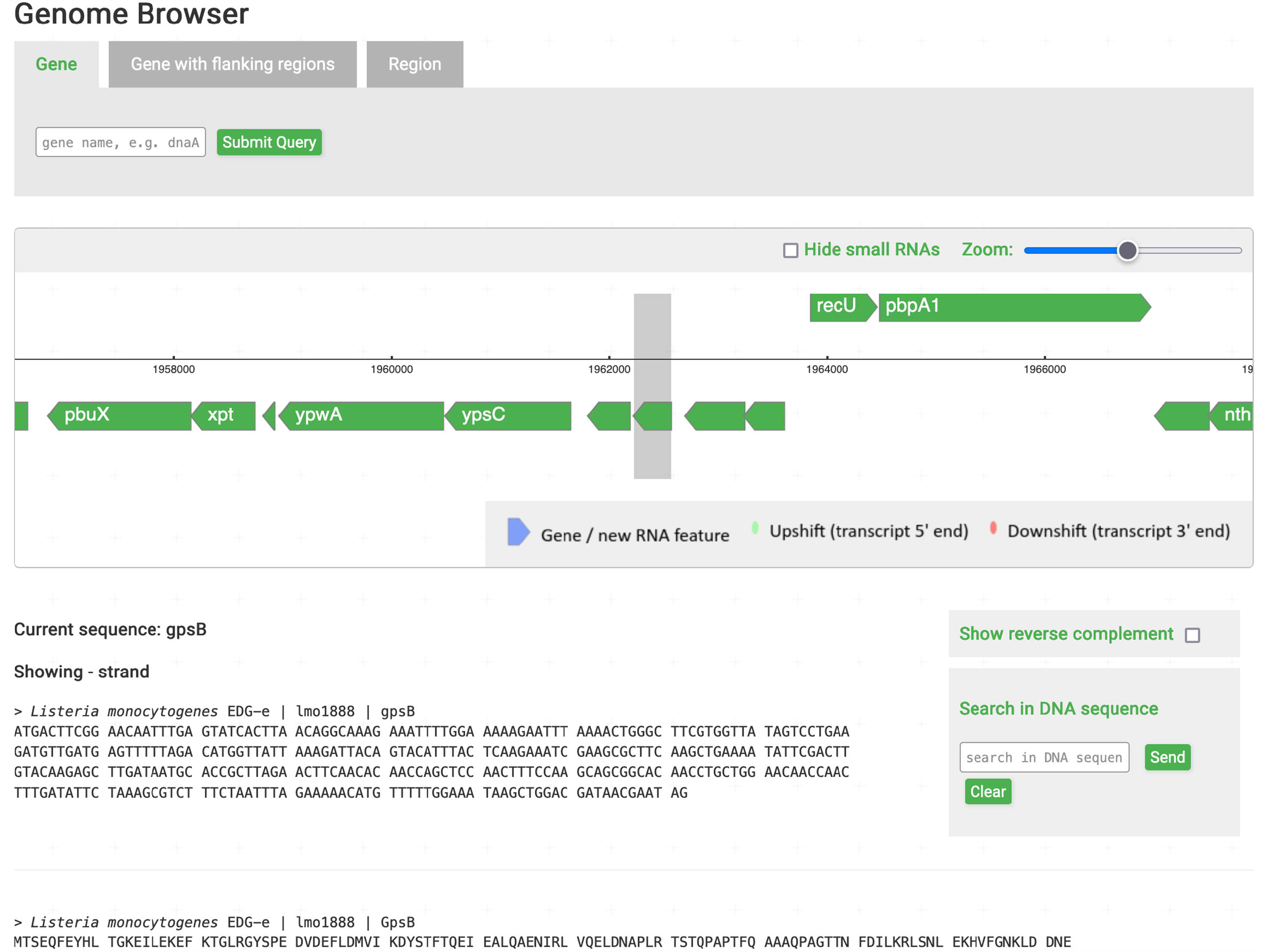
The Genome Browser displaying the genomic context of *gpsB*. The graphical element in the page center allows the user to interactively explore the genome. Below, sequences corresponding to the current selection are shown. The tabs at the page top provide more complex options for searching the genome.

In addition to the aforementioned Interaction Graph, the Interaction Browser (Fig. 5) lets the user explore protein-protein interaction networks in even greater detail. The browser shows a dynamic network of interactions with a range of different options. The user can click on the nodes representing proteins to display more information in the form of a tooltip or drag them around to rearrange them. The settings menu offers several options, such as adjusting the *radius* of the network, which corresponds to the neighborhood depth of the graph. On a low setting, only the selected protein and its direct interaction partner are included. But by increasing the radius, more and more interaction partners of the partners can be included. Another setting, the *spacing*, controls the distance between nodes in the graph. The user can also highlight a protein via a search bar and change the color scheme for nodes and edges. Right-clicking empty space in the interaction display opens a context menu which allows to export the viewed network in different formats. The info box in top left corner contains a search bar as well as the proportion of currently loaded proteins.

**Figure 5.**
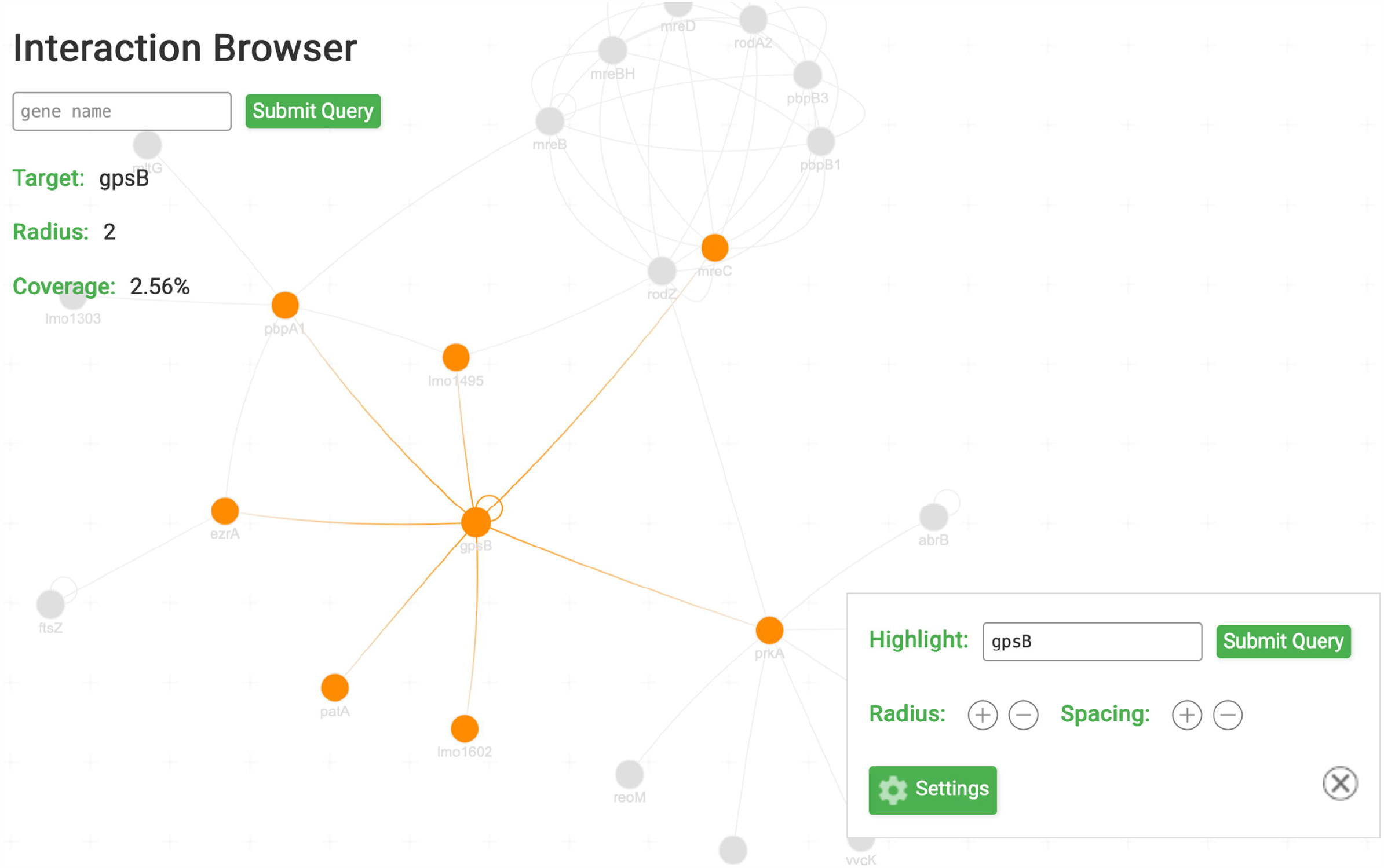
The Interaction Browser with GpsB selected. This interactive tool displays a dynamic network of protein-protein interactions. The user can rearrange the nodes, highlight a specified protein, or increase the radius to include interactions of interaction partners.

The last browser, the Expression Browser (Fig. 6), features a line chart representing the raw intensities of transcript levels for different experimental conditions. The data stems from an analysis of global gene expression in the mouse spleen during infection (Camejo et al., 2009). Using the search bar, the expression of multiple genes can be compared in the same chart. Upon clicking on data points, an explanation of the corresponding experimental condition appears. Data can be exported using the buttons in the top right corner. The plot is drawn using the Apache ECharts visualization library.

**Figure 6.**
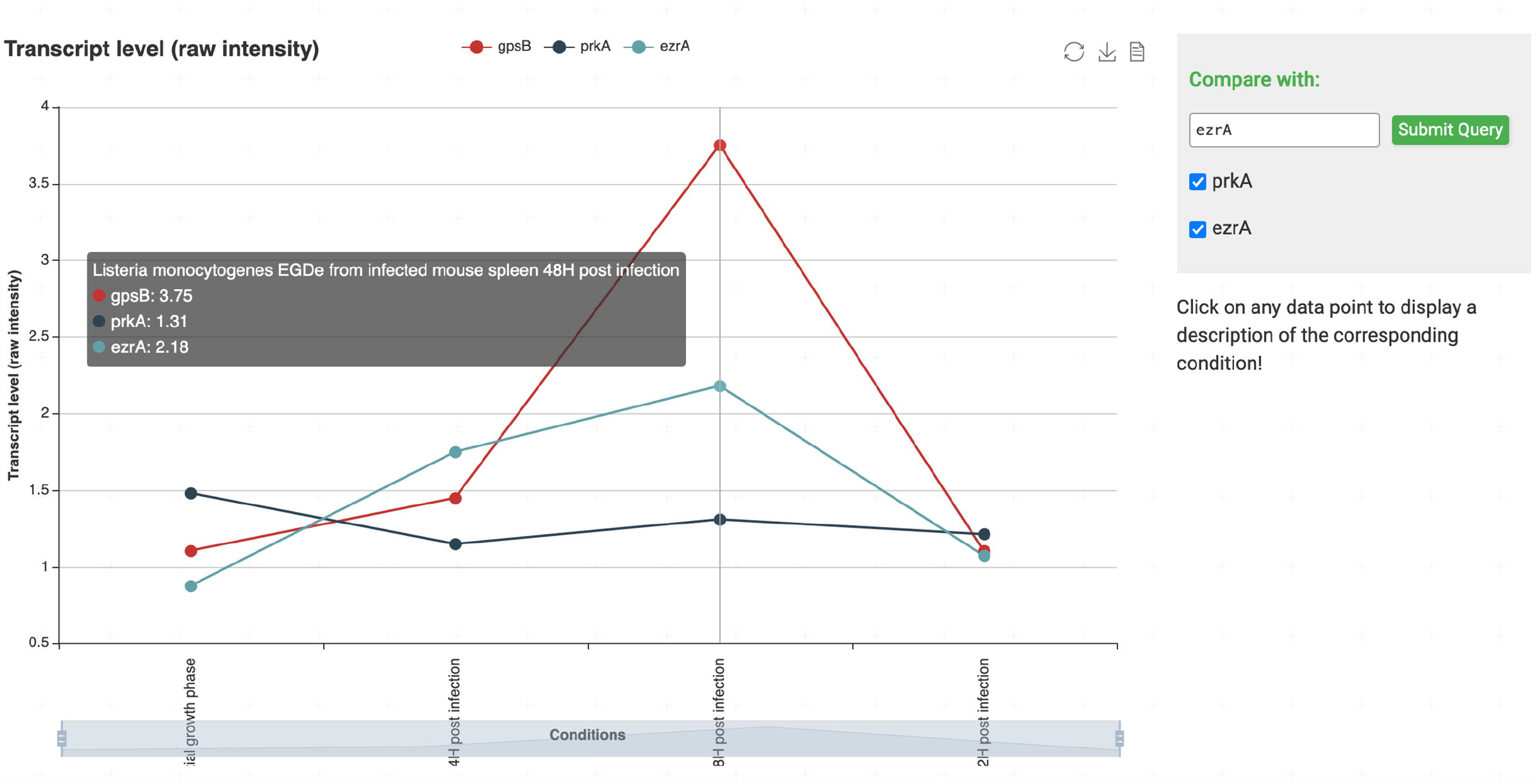
The Expression Browser comparing the transcript levels of *gpsB* (red line), *prkA* (dark blue) and *ezrA* (turqoise) before and during infection. The search bar allows to load data on additional genes into the chart. When hovering over a data point, the corresponding value and description of the experimental condition are displayed in a tooltip. If the user clicks on the data point, the description is also shown below the search bar.

## IMPLEMENTATION AND DATA

*Listi*Wiki uses the same framework as its sibling platforms *Subti*Wiki, *Syn*Wiki and *Myco*Wiki. As a result, the overall design, functionality and organization of data are generally the same. However, availability of certain data and corresponding frontend features might vary between the websites. The backend is written in PHP and hosted with a Apache HTTP server, while the frontend uses simple JavaScript without a specialized framework. The relational database used for data storage is MySQL.

The data featured on *Listi*Wiki is a mixture of manually curated information and bulk imports from other sources. The foundation of data is laid by the genome annotation of *L. monocytogenes* strain EGD-e (RefSeq accession: NC_003210.1) (Toledo-Arana et al., 2009). Gene essentiality and gene expression data were obtained from published information (Camejo et al., 2009; Fischer et al., 2022). Using specific PubMed queries, 1,285 publications were added as relevant references to the Gene pages. Based on homologies to *B. subtilis* genes, the database was populated using *Subti*Wiki as a rich source of information. Besides the aforementioned 328 newly assigned gene names, 1636 protein-protein interactions were adopted from *Subti*Wiki for orthologous proteins.

Similar to its sibling platforms, *Listi*Wiki received a specifically curated set of categories to appropriately group and describe genes in a standardized manner. The categories are structured like a tree, with six top-level categories which are further subdivided for a more fine-grained characterization of genes. For example, genes are grouped by their roles in cellular processes and metabolism, genomic properties, localization, and essentiality, among others. Table 1 gives an overview over the top-level categories and their immediate subcategories.

**Table 1.**
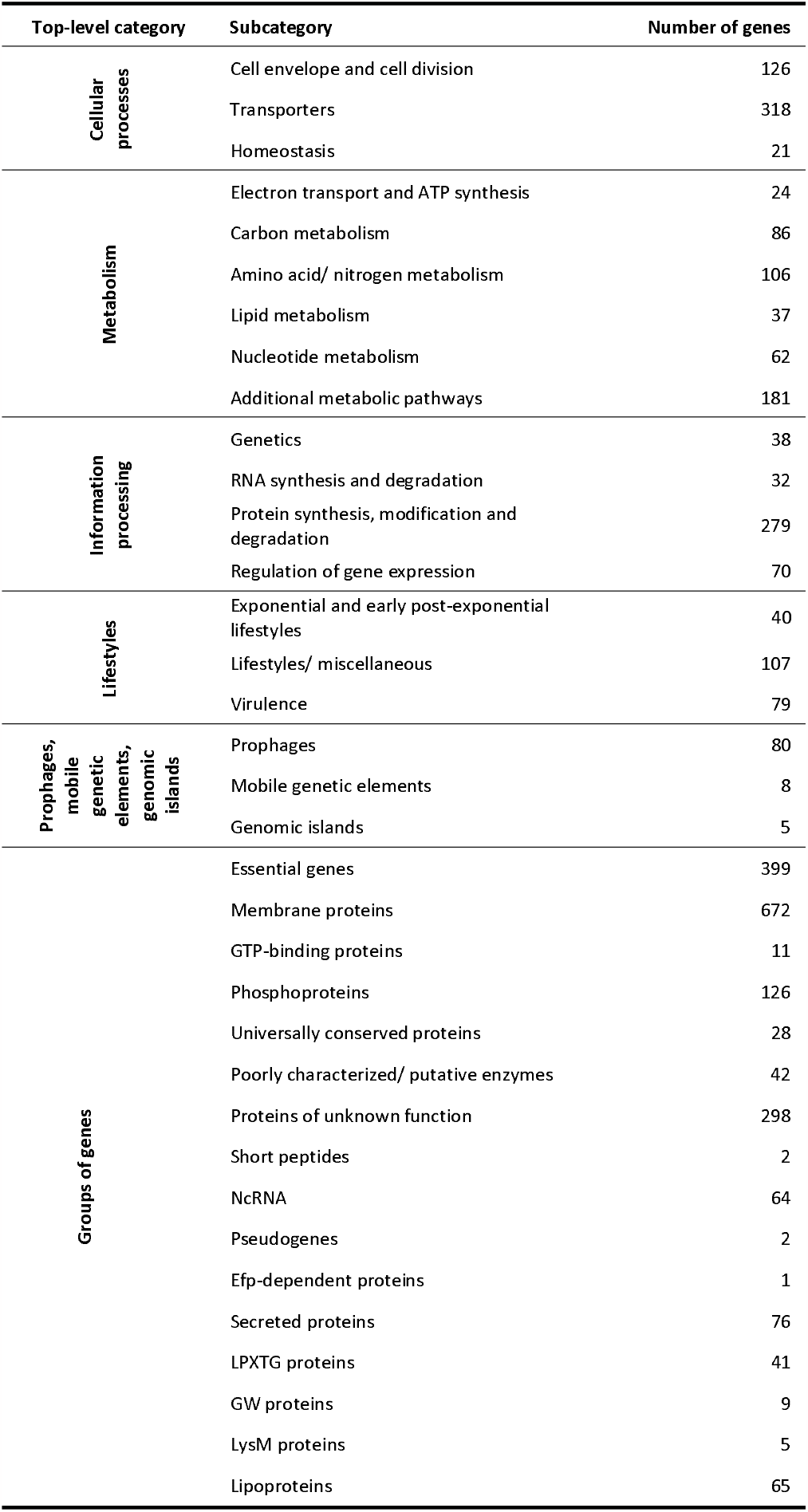
List of top-level categories and their subcategories used in *Listi*Wiki, and the number of genes assigned to each of them.

## CURATION AND COMMUNITY

As keeping up with the newest scientific findings is a continuous effort, we invite the community of *Listeria* researchers to contribute to *Listi*Wiki. It is possible to request a registration using the “Register” button, which can be found on the Front page and on any Gene page. Although exploring the data on *Listi*Wiki does not require any user account, a login is needed for editing the available data and contributing new information. Changes to the database are regularly reviewed to ensure the quality of provided information.

The presentation of up-to-date information and the possibility for the scientific community to contribute to the effort are important advantages of *Listi*Wiki over other genome databases available for *L. monocytogenes. <u>Listi</u>*List as the first of them was published in 2001, along with the announcement of the *L. monocytogenes* EGD-e genome, to facilitate genome handling through a web interface called GenoList (Glaser et al., 2001; Moszer et al., 1995). *Listi*List has been a useful reference for many years and even is still accessible while the service of several other GenoList databases has been terminated. However, the annotation used in *Listi*List has never been updated since its release and also cannot be edited. As a result, the presented information is outdated. Another genome browser has been developed based on the BioCyc platform for the second widely used *L. monocytogenes* reference strain 10403S and a few other *L. monocytogenes* strains. The 10403S BioCyc browser provides additional tools for the visualization of biochemical pathways and transcriptional networks (Orsi et al., 2015). However, the use of the BioCyc database requires a paid subscription, whereas *Listi*Wiki is available to all users without any subscription or charges.

The annotation currently stored in *Listi*Wiki has been compiled from global and gene-specific data sets available in the scientific literature, either by automated or manual import. This initiative needs to be continued and therefore we invite the *L. monocytogenes* scientific community to keep this annotation up to date in a joint effort. *Listi*Wiki is a dynamic database that constantly expands along with the progress of research and which is curated by active scientists in the field. Thus, it may become a useful resource for the easy and barrier-free access of current *L. monocytogenes* functional genome annotation.

## FUTURE PERSPECTIVES

With *Listi*Wiki we have released a new and self-explanatory tool for the intuitive exploration of the genome of this pathogen more than two decades after the release of *Listi*List, the first genome browser that was available for *L. monocytogenes*. By consideration of the gene designations that have been proposed since 2001, *Listi*Wiki translates the results from over 20 years of *L. monocytogenes* research into an updated genome annotation. Moreover, the database features an expanded toolbox with modern functionalities to facilitate the generation of hypotheses on the function and the mutual interactions of genes and their gene products. *Listi*Wiki is a dynamic database and can be complemented and expanded when further data become accessible. Currently, global analyses on protein-protein interactions between *L. monocytogenes* proteins or even on the interactions of *L. monocytogenes* proteins with the host proteome are not available. Likewise, the determination of essential gene sets under varying conditions has just started recently (Fischer et al., 2022; Wu et al., 2022). When information on gene essentiality in different mutant backgrounds becomes available one day, implementation and visualization of such synthetic lethal interactions could possibly be a useful extension. Besides this, *Listi*Wiki represents the fourth member of a family of similar databases that includes *Myco*Wiki, *Subti*Wiki, and *Syn*Wiki. Thus, *Listi*Wiki will profit from the introduction of novel features that will be developed for these other three databases.

## ACKNOWLEDGEMENTS

We are grateful to Marie-Theres Thieme, Fabian Commichau, Richard Lüdtke and Johannes Gibhardt for their help with the initial work on *Listi*Wiki. JS and SH would like to thank the German Research Foundation for the longstanding financial support of the research in their laboratories.

## Notes

### Competing Interest Statement

The authors have declared no competing interest.

### Summary of Updates

a typo in the title was corrected

